# Cross-fostering in rodents causes region-specific alterations in entorhinal cortical gamma rhythms associated with NMDA receptor dysfunction

**DOI:** 10.1101/298612

**Authors:** Stephen Hall, Karen Hawkins, Grace Laws, Thomas Akitt, Anna Simon, Ceri H. Davies, Miles A. Whittington, Mark O. Cunningham

**Author notes:** Author for correspondence: Mark Cunningham, Institute of Neuroscience, The Medical School, Newcastle University, Framlington Place, Newcastle upon Tyne, NE2 4HH, UK.

## Abstract

There has recently been a large increase in the number of children placed in foster care in the United States and Europe. While this is ‘the least worst scenario’ for those with a lack of appropriate biological care, it is recognised that these children are exposed to major stressors correlated with behavioural changes, particularly in the realm of social cognition into adulthood. Here we model foster care in rodents: rat pups are removed from their biological mother and placed with a non-genetically related dam. This prevented the entorhinal cortex from generating patterns of gamma rhythms required for normal parahippocampal function relevant to social interaction. These changes correlated with a reduction in NMDA receptor-mediated excitation, and changes in parvalbumin expression in interneurons. These data suggest that early life care delivered by a non-biological parent may disrupt social behaviour but, in contrast, generate neurobiological changes antagonistic to those currently associated with psychosis.

**Significance Statement:** Cross fostering is an effective approach for delineating the effect of environment from genetic influences upon behavior. This involves removal of pups from one mother and transfer to another lactating dam. This manipulation is considered as a mild form of early life stress, producing neurobehavioral changes such as alterations in social interaction. We demonstrate that cross fostering produces changes in the ability of cortical microcircuits to generate oscillatory rhythms, in particular the gamma rhythm, in brain regions important for social cognition. This reduction in gamma rhythmogenesis is related to a reduction in synaptic drive provided by the NMDA receptor. One implication of this work is that the modulation of NMDA receptors offers a potential therapeutic strategy for disorders involving impaired sociability.

## Introduction

An increasing number of people spend their childhood and adolescence separated from their biological parents. Currently in the UK 6 in 1000 under 18 year olds are looked-after by local authorities, with ca. 80% of them in foster care^1^. The most recent figures from the US^2^ show that approximately 641000 children have spent time in foster care and that a significant proportion have experience deprived and chaotic environments for a period of time^3^. While this situation represents the optimal scenario for children with absent or inappropriate biological parenting, it has long been postulated that the care a child receives during their early years could have a significant impact upon their mental health in later life^4^. Recently it has been shown that children who suffer poor maternal care can develop behavioural and neurodevelopment problems^5–7^ and further to this, children who experience long-term foster or institutional care have a higher likelihood of social, behavioural and neurological problems^8,9^. Children with any history of institutionalization have more psychiatric disorders than those without at 54 months of age^10^. However, children placed into foster care exhibit less internalizing symptoms, than those remaining in institutional care^10^. The follow up of this study, when the children were aged 12 years old, showed that children placed in foster care had fewer externalizing symptoms than those receiving institutionalized care, but the counts for internalizing symptoms and ADHD remained the same^11^. Furthermore, unemotional-callous behavior, a potential precursor to psychopathy, was significantly higher in institutionalized children, as compared to non-institutionalized controls^11^. These studies suggest that the quality of foster care could have a significant impact upon the outcomes for children reaching into their adulthood.

A range of early life manipulations have attempted to replicate poor maternal care to permit more mechanistic studies in experimental animal models. Neonatal handling, where pups are removed from their mother and handled for a set period of time, has shown permanent increases in glucocorticoid receptor density^12^ and increases in the NR2B subunit of NMDA receptors in the hippocampus^13^. Isolation rearing, whereby offspring are separated from littermates after weaning, shows changes in both the function^14^ and the structure^15,16^ of cells within the parahippocampal region. The most extensively researched early life manipulation, maternal separation, in which pups are removed then returned to the mother for set periods during weaning, show reductions in hippocampal cell neurogenesis and structural reorganisation of the dentate gyrus^17^. Other studies have also shown neurodevelopmental changes within the hippocampus^18^. These studies suggest that the parahippocampal region exhibits significant changes following early life manipulation which may lead to altered function into adulthood.

In the above models, rodent pups undergoing maternal separation or maternal deprivation paradigms mimic a range of neurodevelopmental problems seen in some children in care^19–21^. However, these animal models are severe. We argue that a more accurate way to mimic the social situation and to examine the core stressors associated with foster care in humans is cross-fostering: Cross-fostering replaces the mother of the offspring with a non-biological mother. While no maternal deprivation occurs *per se*, research shows that behavioural consequences are much more akin to those seen in studies of human foster care. For example, cross-fostered macaques show less vocalisation and less social interaction than those remaining with their birth mothers^22^ and cross-fostered mice exhibit reduced social interaction times and a decrease in social interaction when compared to the control group^23^. These authors also showed that cross-fostering brought about changes in emotional behaviours such as decreased exploratory behaviour and an increased stress response.

We therefore used the cross-fostering model in rats to examine possible mechanisms underlying the abnormal development of social behaviours observed in adults placed in care during childhood. The parahippocampal region has been strongly indicated in more severe animal models of early life stress (above), but has been underexplored in cross-fostering. As such we focussed on gamma oscillation generation in the medial entorhinal cortex (mEC) and the CA3 region of the hippocampus. Changes in gamma oscillations have been observed in many models of cognitive and behavioural deficits in rodents^24,25^ and here we show that cross-fostering produces an alteration in gamma rhythmogenesis predominantly in mEC which is associated with altered NMDA receptor function and parvalbumin-immunopositive interneuron number.

## Results

### Cross fostering caused fragmentation of post-natal maternal care

During the course of the first six days following birth naturally reared (N) litters were predominantly restricted to one nest by the dam. In contrast, cross fostered (CF) litters were split into multiple nests (N, 1.0 ± 0.02 vs CF, 1.5 ± 0.05, P<0.05, n=4 for N and 3 for CF, Fig. 1A). Analysis of the time pups spent away from the nest also revealed significant differences between N and CF groups. The average duration of time spent out of the nest over the first six days was significantly greater in CF pups than N pups (N, 121.2 ± 32.1secs vs CF, 249.9 ± 38.5 secs; P<0.05, n= 4 for N and 3 for CF, Fig. 1B). In contrast, there was no significant effect of CF upon maternal nursing behaviours such as licking-grooming (LG), physical contact, arched back nursing (ABN) and passive nursing (P>0.05, n=4 for N and 3 for CF, Fig. 1C).

**Figure 1.**
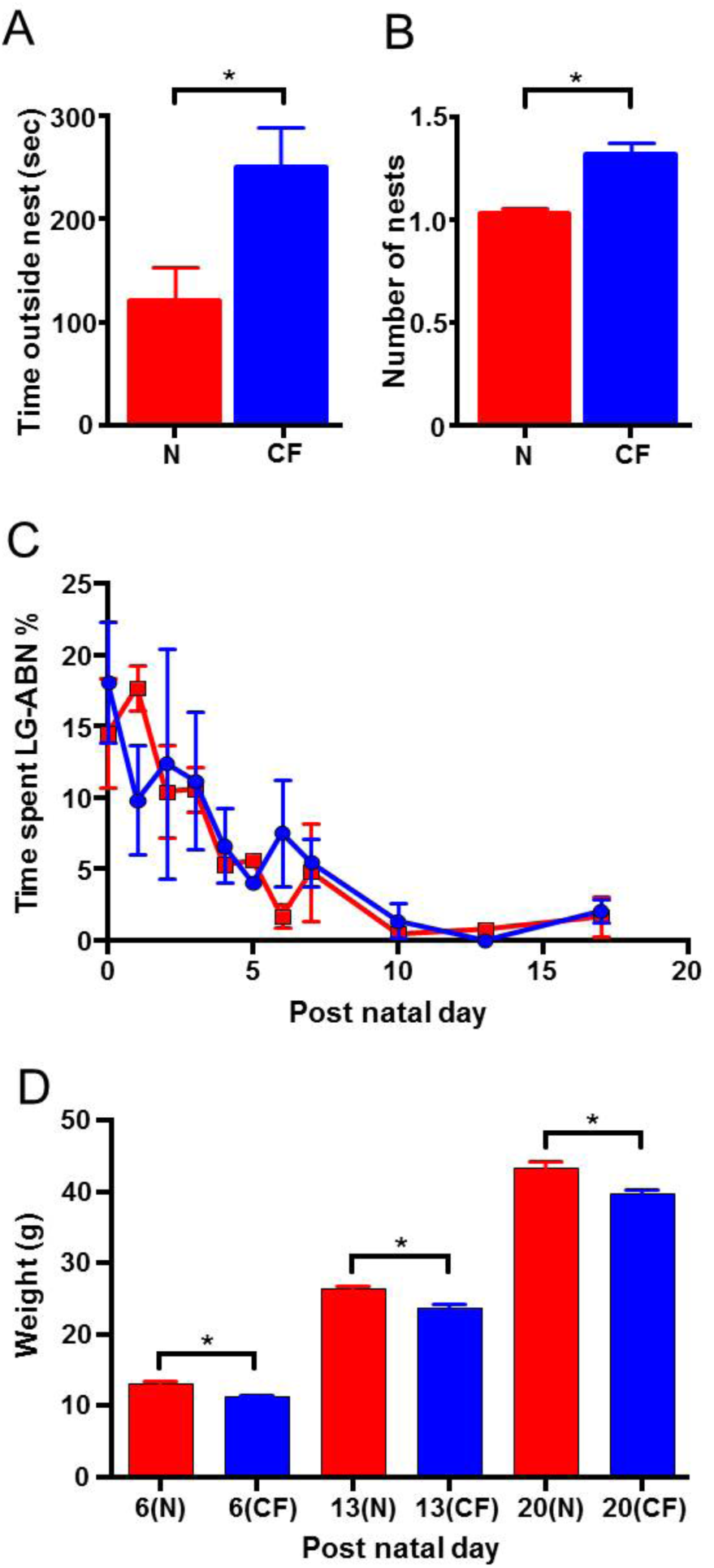
Cross-fostered pups experience fragmented maternal care as compared to naturally reared offspring. (A) Average duration of time spent by pups outside of the nest over the first five days following birth. (B) Average number of nests for N and CF litters across the first five days following birth. (C) Mean weights of CF and N pups across the pre-weaning period. (D) Time spent investing in the licking grooming and arch backed nursing (LG-ABN) behaviour in the pre-weaning period on PND 0-7, 10, 13 and 17. Data is represented as mean percentage ± SEM of the 1000 seconds of behaviour analysed per day per animal.

These alterations in the dam and pup behaviour were associated with a significant difference in the body weight between (CF) and N groups. CF pups exhibited a small but significantly lower rate of body weight gain during development (Fig. 1D) across all the age time-points assessed (PND6: N, 13.1 ± 0.3g vs CF, 11.2 ± 0.2 g, P<0.005. PND13: N, 26.4 ± 0.5g vs CF, 23.8 ± 0.4 g, P<0.05. PND 20: N, 43.4 ± 0.9g vs 39.7 ± 0.6 g, P<0.05, n=7 for N and 8 for CF across all PND values).

### Cross fostering caused greater alterations of gamma oscillations in mEC compared to hippocampus in adults

The hippocampus and mEC are potent generators of gamma frequency oscillations in adult rats both *in vivo*^26,27^ and *in vitro*^28–31^. When increasing concentrations of kainate (KA) were added to hippocampal CA3 in slices from N and CF animals (Fig. 2Ai,ii, Ci) no significant change in peak frequency was observed across the entire KA concentration range used (n=8 slices, 6 animals, P>0.05). There was a trend towards a decreased frequency in CA3 slices from CF rats through a concentration range of 1-500 nM KA; for example, CA3 from N rats: 1 nM KA, 32.6 ± 3.5 Hz; 20 nM KA 41.4 ± 2.1 Hz), CA3 from CF rats: 1 nM KA, 26.6 ± 3.1 Hz; 20 nM KA, 35.2 ± 3.9 Hz. However no overall significant difference was seen between gamma frequency generated in slices from N and CF rats (*P*>0.1).

**Figure 2.**
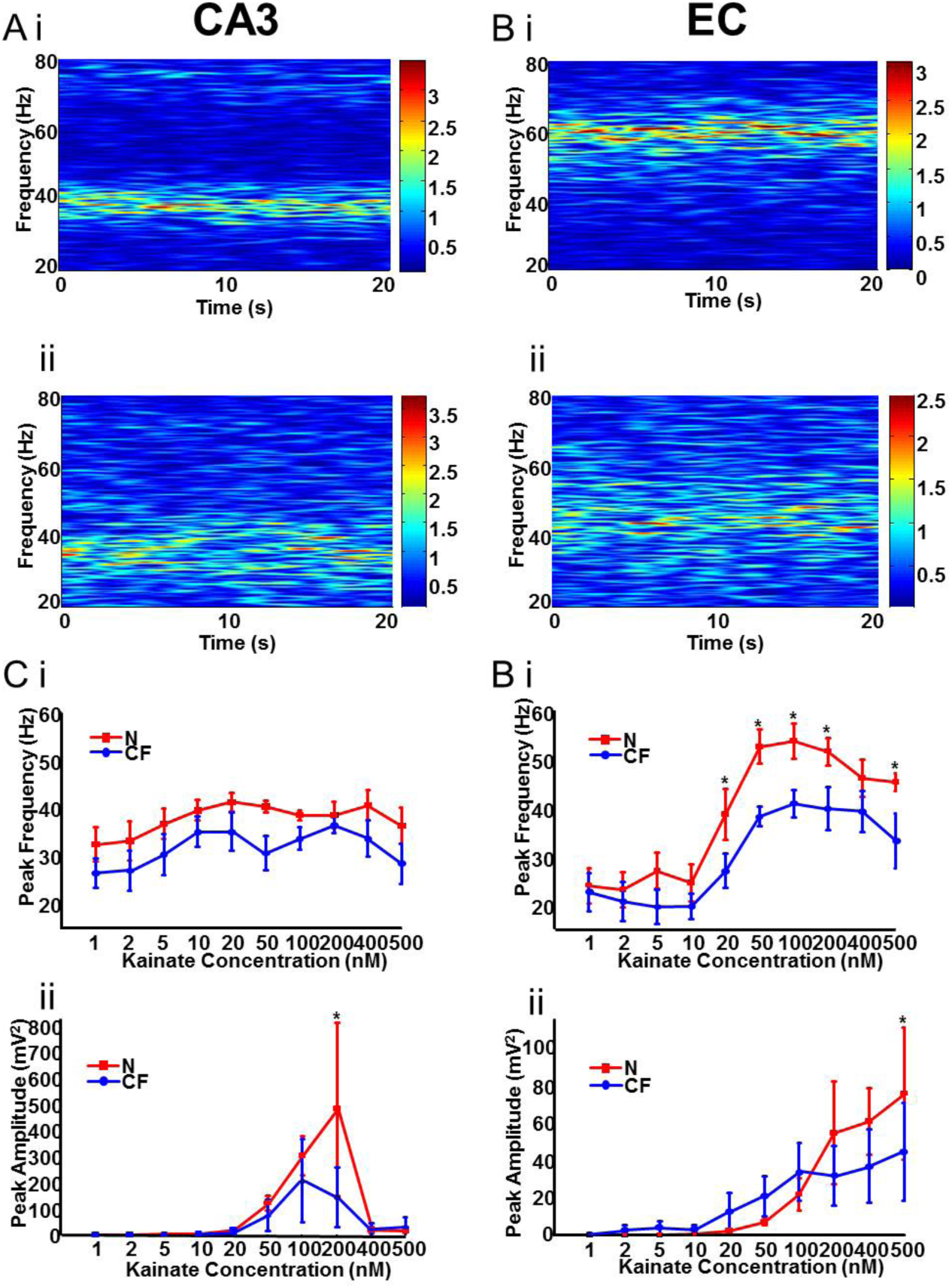
Increasing concentrations of KA revealed peak power and frequency differences between natural and cross-fostered rodents. (A) Example spectrograms generated from a 20s epoch of CA3 local field potential recordings in the presence of 100nM kainate in both N (i) and CF (ii) slices. (B) Example spectrograms generated from a 20s epoch of mEC local field potential recordings in the presence of 100nM kainate in both N (i) and CF (ii) slices. (C) (i) Graph showing the peak frequency of gamma oscillatory activity in CA3 with increasing concentrations of KA. (ii) Graph showing the peak power of gamma oscillatory activity in CA3 with increasing concentrations of KA. (D)(i) Graph showing the peak frequency of gamma oscillatory activity in mEC with increasing concentrations of KA. (ii) Graph showing the peak power of gamma oscillatory activity in mEC with increasing concentrations of KA.

In contrast, when increasing concentrations of KA were added to the mEC of brain slices obtained from N and CF animals (Fig. 2Bi, ii, Di), the frequency of the oscillations showed a sharp, concentration-dependent increase (n=9 slices, 6 animals for each group). At concentrations from 1-10 nM, KA-induced oscillations were relatively similar in slices from both N and CF rodents, with both showing beta/low-gamma frequency oscillations (1-10nM average: N, 24.7 ± 1.6 Hz; CF 20.7 ± 1.5 Hz). At concentrations of 20 nM or higher, gamma oscillation in EC from CF rats failed to increase as much as those in EC from N rats. At 100 nM kainate gamma rhythms reached 56.0 ± 2.9 Hz in EC from N rats but only 41.1 ± 2.2 Hz in EC from CF rats (P<0.05). This significant reduction in peak frequency was seen from 20-200 nM kainate and at 500nM kainate (Fig. 2Di).

Very little change in kainate-induced gamma power was seen either in CA3 (Fig. 2Cii) or EC (Fig. 2Dii) when considering the entire concentration range of KA (P>0.05). However, the peak powers seen in CA3 and EC from N rats (at 200 and 500 nM KA respectively) were significantly larger than those seen in slices from CF rats at these concentrations (P<0.05 in each case). In addition, using cholinergic receptor activation with carbachol (5 μM) to generate gamma oscillations^29,30^ failed to generate significant changes in gamma frequency in either region when comparing slices from N and CF rats. In both regions, there was no change in peak gamma frequency (EC N; 33.3 ± 1.6 Hz vs. CF; 32.9 ± 2.9 Hz, CA3 N; 30.1 ± 3.9 Hz vs. CF; 30.5 ± 5.0 Hz,) or power (EC N; 38.4 ± 25.5 Hz vs. 62.2 ± 39.4 Hz, CA3 421.7 ± 254.1 μV^2^ vs. 527.2 ± 376.7 μV^2^. n = 9 slices, 5 animals, P > 0.05 for all, data not illustrated).

### NMDA receptor signalling was involved in the EC-specific changes in gamma frequency and amplitude in CF rats

A similar decrease in the above KA-induced gamma oscillation frequency and power in EC from CF rats has been seen previously in control rats by application of ketamine^31^. Acute ketamine administration generates social withdrawal and psychosis-like effects in subjects^33^ which may be related to the report of increased incidence of social problems and psychosis in adults previously in care^10^. To test whether the changes seen in CF rats correlated with ketamine-like effects an occlusion experiment was performed. Ketamine (25 μM) was perfused prior to adding increasing concentrations of KA to the EC or hippocampus of N or CF rodents.

In slices obtained from the CA3 of N rodents, ketamine caused a small but significant decrease in gamma frequency at kainate concentrations above 100 nM (Fig. 3A, P<0.05). No such change in gamma frequency was seen for kainate-induced gamma rhythms in CA3 from CF rodents (*P*>0.05 for all concentrations). Similarly, in slices obtained from the EC of N rodents, at concentrations of 20 nM kainate and upwards, ketamine caused significant decreases in peak gamma frequency (n=7 slices, 4 animals, P<0.01, Fig. 3B). No significant changes in EC gamma frequency, caused by ketamine, were seen in slices from CF rodents (n=7 slices, 4 animals, P>0.05).

**Figure 3.**
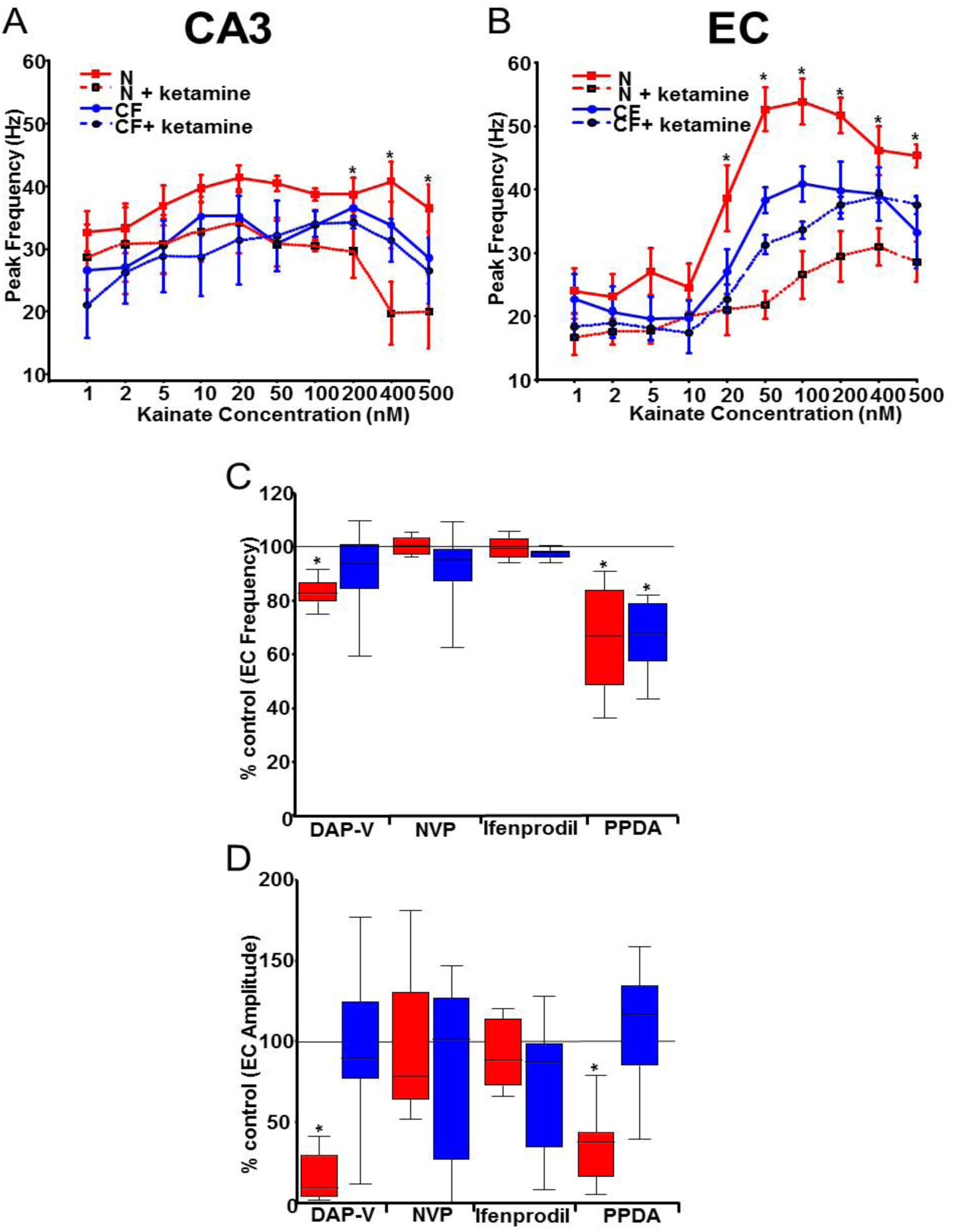
Slices from natural and cross-fostered rats showed different responses to NMDA receptor antagonism. (A) Graph showing the peak frequency of gamma oscillatory activity in CA3 against concentration of KA. Dashed lines show the peak frequency after NMDA receptor antagonism with 25μM ketamine. (B) Graph showing the peak frequency of gamma oscillatory activity in the EC against concentration of kainate. Dashed lines show the peak frequency after NMDA receptor antagonism with 25μM ketamine. (C) Box plot showing the % change in peak gamma frequency after the application of NMDA receptor antagonists. Red boxes = N, blue boxes = CF. (D) Box plot showing the % change in peak gamma power after the application of NMDA receptor antagonists. Red boxes = N, blue boxes = CF.

Ketamine is a broad-spectrum NMDA receptor antagonist with additional effects on a number of other neurotransmitter systems. We therefore explored the nature of this apparent occlusion of ketamine’s effects in slices from CF rats in more detail. We focused on the EC, where the effects of cross-fostering and ketamine were greatest cf. CA3. Application of D-AP5 (50 μM), a more selective antagonist for NMDA receptors than ketamine, to gamma oscillations in EC slices obtained from N rodents significantly reduced the peak frequency from 47.6 ± 1.8 Hz to 38.6 ± 3.2 Hz and peak power from 84.7 ± 35.8 μV^2^ to 4.7 ± 0.9 μV^2^ (n=11 slices, 5 animals, P<0.01; Fig. 3C,D). In contrast, in EC slices obtained from CF rodents D-AP5 caused no significant changes (P>0.05, n=9 slices, 5 animals; Fig. 3C,D) in either gamma frequency (46.8 ± 3.1 Hz to 44.6 ± 2.8 Hz) or gamma peak power (101.2 ± 68 μV^2^ to 86.5 ± 45.3 μV^2^).

More selective antagonism aimed at NR2A subunit-containing NMDA receptors with application of NVP (50nM) caused no significant change in the peak power (N; 32.8 ± 6.7 μV^2^ vs 30.3 ± 5.03 μV^2^. CF; 32.3 ± 10.3 μV^2^ vs 25.9 ± 11.25 μV^2^) or the peak frequency (N; 43.8 ± 1.7 Hz vs 43.3 ± 1.9 Hz. CF; 45.8 ± 2.2 Hz vs 45.0 ± 3.1 Hz) of gamma oscillations in both N (n=8 slices, 4 animals, P>0.05) and CF slices (n=9 slices, 4 animals P>0.05). Similarly, targetting the antagonism of NR2B-containing NMDA receptors via application of Ifenprodil (10μM) caused no significant changes in peak power (133.34 ± 117.2 μV^2^ vs 119.2 ± 103.9 μV^2^, n=9 slices, 4 animals, P<0.005) or gamma frequency (45.1 ± 1.5 Hz vs 45.1 ± 1.4 Hz, n=9 slices, 4 animals P>0.05) in N slices. In cross-fostered animals, antagonising the subunit also caused no significant difference in the peak frequency (46.1 ± 1.0 Hz vs 44.7 ± 0.9 Hz) or peak power (117.4 ± 89.2 μV^2^ vs 104.9 ± 83.8 μV^2^) of gamma oscillations (both n=8 slices, n=4 animals, P>0.05).

In contrast, antagonism aimed preferentially at GluN2C/D subunit-containing NMDA receptors with PPDA (100nM) caused large reductions in both peak frequency (43.6 ± 2.1 Hz vs 31.8 ± 3.4 Hz) and peak power (48.3 ± 15.9 μV^2^ vs 10.3 ± 3.7 μV^2^) in EC slices from N animals (n=7 slices, 5 animals, P<0.005). In EC slices from CF rats, antagonising the subunit also caused a significant reduction the peak frequency of gamma oscillations (46.1 ± 1.1 Hz vs 34.2 ± 2.4Hz, n=7 slices, 4 animals, P<0.05, Fig. 3C). No significant change in the peak power of gamma oscillations was seen in EC from CF rats (45.1 ± 16.9 μV^2^ vs 55.8 ± 27.1 μV^2^ n=7 slices, 4 animals, P>0.05).

### Cross fostering had only minor effects on synaptic inhibition onto EC principal cells

The above experiments strongly suggested that a reduced functionality in NMDA receptors (predominantly GluN1-GluN2C/D containing subtypes preferred by ketamine) may underlie the changes in gamma rhythm generation seen in EC of CF rats. Unlike many neocortical areas, this region retains a large NMDA receptor-mediated excitation of interneurons in adulthood^34^ and loss of this drive to basket cells has been shown to reduce power and frequency of gamma rhythms^32^. However, decreased gamma frequencies can also be generated by changes in the size and decay kinetics of GABA_A_ receptor-mediated synaptic inhibition35,36. We therefore examined the profile of IPSPs during gamma rhythms and evoked by electrical stimulation to quantify any changes in this form of inhibition to principal cells.

During kainate evoked population gamma activity recordings from both N and CF slices in layer III pyramidal neurons (Fig. 4Bi) and layer II stellate cells (Fig. 4 Bii) revealed rhythmic IPSPs that were phase locked to the local field potential rhythm. During gamma oscillatory activity, recordings from layer III pyramidal neurons in CF slices revealed IPSP trains with peak power (0.025 ± 0.014 mV^2^) significantly larger than the power of IPSP trains during gamma rhythms in EC slices from N rats (0.013 ± 0.001 mV^2^) (*P*<0.05, n=9 cells each group, 9 rats). The fundamental frequency of these trains of IPSPs was slightly lower in CF pyramidal neurons (50.7 ± 4.2 Hz) compared to N pyramidal neurons (57.9 ± 9.5 Hz) but this finding was not significant (*P*>0.05). In stellate cells, no significant difference in mean IPSP power was seen during gamma rhythms (Fig. 4 Bii). The peak power of IPSP trains in N stellate cells (0.023 ± 0.013 mV^2^) compared to CF stellate cells (0.012 ± 0.014 mV^2^) (*P*>0.05, n=11 cells each group, 11 rats). As with IPSP trains in pyramidal cells, the frequency of the IPSP trains in stellate cells was also not significantly altered (*P*>0.05).

**Figure 4.**
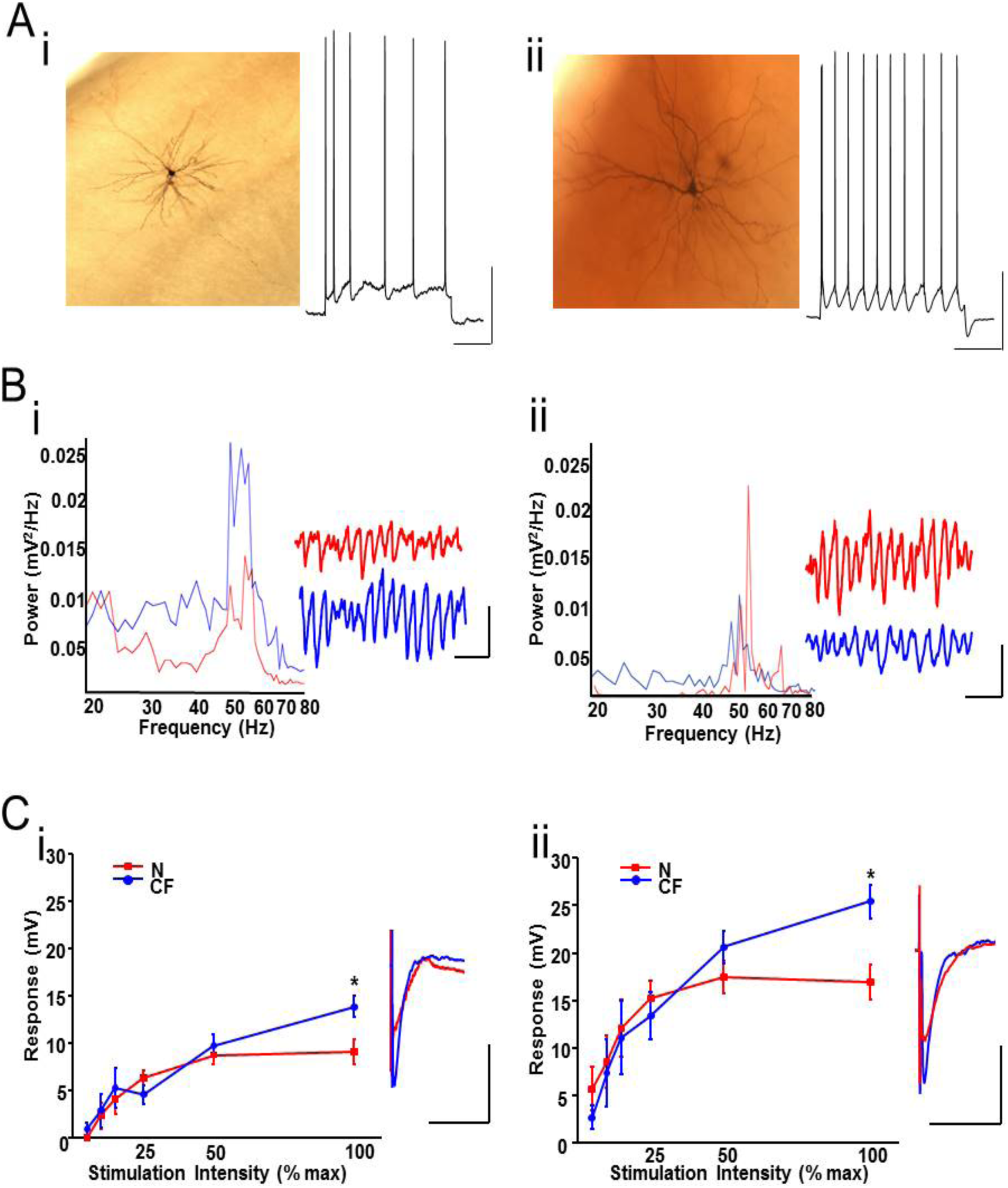
Cross fostering generated subtle differences in inhibitory synaptic input onto principal cells. (A) Biocytin reconstruction of a pyramidal cell (i) and a stellate cell (ii) from the superficial layers of the mEC and response to step depolarisation. Scalebars 20mV, 100ms. (B) Pooled power spectra of IPSPs from pyramidal cells (i) and stellate cells (ii) during gamma activity evoked by KA in slices from N (red) and CF (blue) rats. Example traces show IPSP trains during gamma rhythms in each case. Scale bars 3mV, 50ms. (C) Graphs showing the amplitude of electrically-evoked monosynaptic IPSP responses in pyramidal cells (i) and stellate cells (ii) from N (red) and CF (blue) slices.

The observed decrease in mean IPSP amplitude during gamma rhythms seen in pyramidal cells could have been generated by a decrease in excitation of presynaptic interneurons (as suggested by the data summarized in Fig. 3) or by either a decrease in GABA release presynaptically and/or a change in postsynaptic receptor density. To separate these two putative mechanisms, we switched to recording monosynaptic IPSPs evoked by electrical stimulation directly in stellate cells and pyramidal neurons under non-oscillating conditions in which GABA_B_ receptor-mediated and ionotropic glutamate receptor-mediated events were blocked pharmacologically.

The magnitude of evoked monosynaptic IPSPs in pyramidal cells was greater in slices from CF rats only at the highest stimulus intensity used (CF 14.0±1.1mV, N 9.1±1.3mV, *P*<0.05, n=12 cells, 9 animals for N and n=9 cells, 8 animals for CF, Fig 4Ci). At this highest level of electrical stimulation, IPSPs in slices from CF rats also had significantly faster kinetics (τd 17.1 ± 2.9 ms in CF vs. 22.7 ± 1.4 ms in N, *P*<0.05). Monosynaptic IPSPs recorded in stellate cells were also larger in CF animals only at the maximal stimulus intensity used in the present study (25.4 ± 1.7mV) compared to N animals (16.8 ± 1.9mV) (*P*<0.05, n=12 cells, 10 animals for N and n=10 cells, 7 animals for CF, Fig. 4Cii). At this stimulation intensity, evoked IPSPs were also significantly faster (τd 20.0 ± 2.7 msecs), than in N animals (τd 34.4 ± 10.9msecs), (P<0.05).

### Cross fostering elevated parvalbumin immunopositive cell count selectively in EC LIII

The NMDA receptor antagonism and IPSP amplitude and kinetic data above strongly suggested that the reduced gamma frequencies seen in EC in slices from CF rats were primarily a consequence of reduced NMDA receptor-mediated excitation to interneurons involved in gamma rhythm generation (mainly parvalbumin (PV) immunopositive basket cells^37^. This interneuron subtype changes its PV expression levels according to how much NMDA receptor-mediated excitation they receive (e.g. see Kinney et al.,^38^) and absence of PV expression has been shown to hugely elevate gamma power, but not frequency^39^. We therefore examined any correlations between the decreased gamma power and frequency seen in CF rats and PV expression levels.

Post hoc immunohistochemical analysis of parvalbumin expression levels in EC slices from CF rats showed a significant increase in positive cells in both layer 3 (LIII) and deeper layers (LV/VI) (P<0.05, n=12 slices, 5 animals for both N and CF, Fig. 5A). The number of PV+ cells in LIII of the mEC was 1708 ± 222 in CF slices compared to 962 ± 114 in N slices, and in LV/VI 1828 ± 179 in CF slices compared to 1330 ± 192 in N slices. No significant changes were observed between N and CF slices in LII (822 ± 119 in N slices vs 741 ± 85 in CF slices, P>0.05, n=12 slices, 5 animals for both N and CF). Staining for GAD67 (GABAergic) neurons, showed no significant differences (P<0.05, n=5 slices, 5 animals for both N and CF) between positive cell number in N or CF slices in either LII (N 805 ± 191 vs CF 756 ± 118), LIII (N 1516 ± 238 vs CF 1820 ± 98) or LV/VI (N 2030 ± 322 vs CF 1764 ± 185). Staining for calretinin containing neurons (expressed only in a few percent of gamma-generating, PV+ basket cells) also showed no significant differences in cell number between slices from N and CF rats (P<0.05, n=5 slices, 5 animals for both N and CF): LII (N 840 ± 121 vs CF 980 ± 149), LIII (N 1003 ± 107 vs CF 1192 ± 170), LV/VI (N 1680 ± 259 vs CF 2100 ± 204). Staining for Somatostatin immunopositive neurons also showed no significant differences (P<0.05, n=5 slices, 5 animals for both N and CF) between detected cell number in N or CF slices in either LII (N 700 ± 87 vs CF 630 ± 102), LIII (N 910 ± 121 vs CF 875 ± 166) or LV/VI (N 1120 ± 175 vs CF 1190 ± 109).

**Figure 5.**
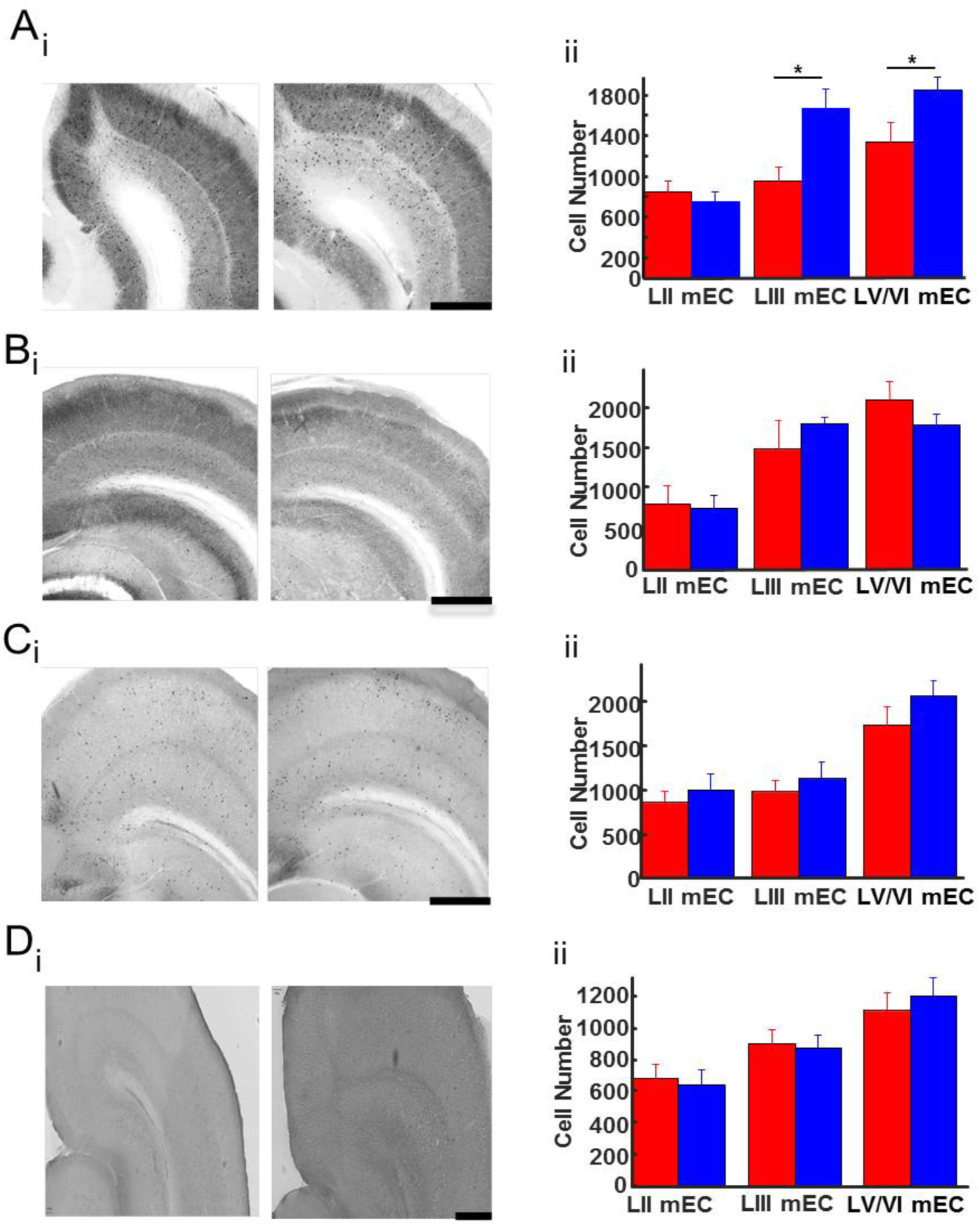
Interneuron expression is altered in slices from cross-fostered rodents, compared to natural rodents. (A) (i) Picture of parvalbumin staining in N (left) and CF (right) sections. (ii) Graph showing the expression of PV^+ve^ interneurons observed in each layer of the EC. (B) i) Picture of GAD67 staining in N (left) and CF (right) sections. (ii) Graph showing the expression of GAD67+ve interneurons observed in each layer of the EC.(C) i) Picture of calretinin staining in N (left) and CF (right) sections. (ii) Graph showing the expression of Cal^+ve^ interneurons observed in each layer of the EC. (D) (i) Picture of somatostatin staining in N (left) and CF (right) sections. (ii) Graph showing the expression of SOM^+ve^ interneurons observed in each layer of the EC.

### Changes in EC gamma rhythms were more overt in ventral rather than dorsal EC

Differences between the ventral and dorsal areas of the parahippocampal region have been highlighted at the level of synaptic inhibition and NMDA receptor-mediated excitation^40^. Furthermore, it has been shown that memory relevant to social interaction (disrupted in adults brought up in care (see introduction)) is selectively associated with the ventral part of the parahippocampal region^41^. Given these precedents, we examined the locus of the observed changes in mEC gamma frequency in more detail. We used a sagittal slice preparation of the mEC to facilitate simultaneous measurement of gamma rhythms in LIII of both dorsal and ventral aspects of mEC using 200 nM KA (Fig. 6). In slices from N animals there was a significant difference in peak gamma frequency between the dorsal (35.9 ± 3.1 Hz) and ventral (62.7 ± 1.2 Hz) mEC regions (P<0.05, n=17 slices, 6 animals, Fig. 6C).

**Figure 6.**
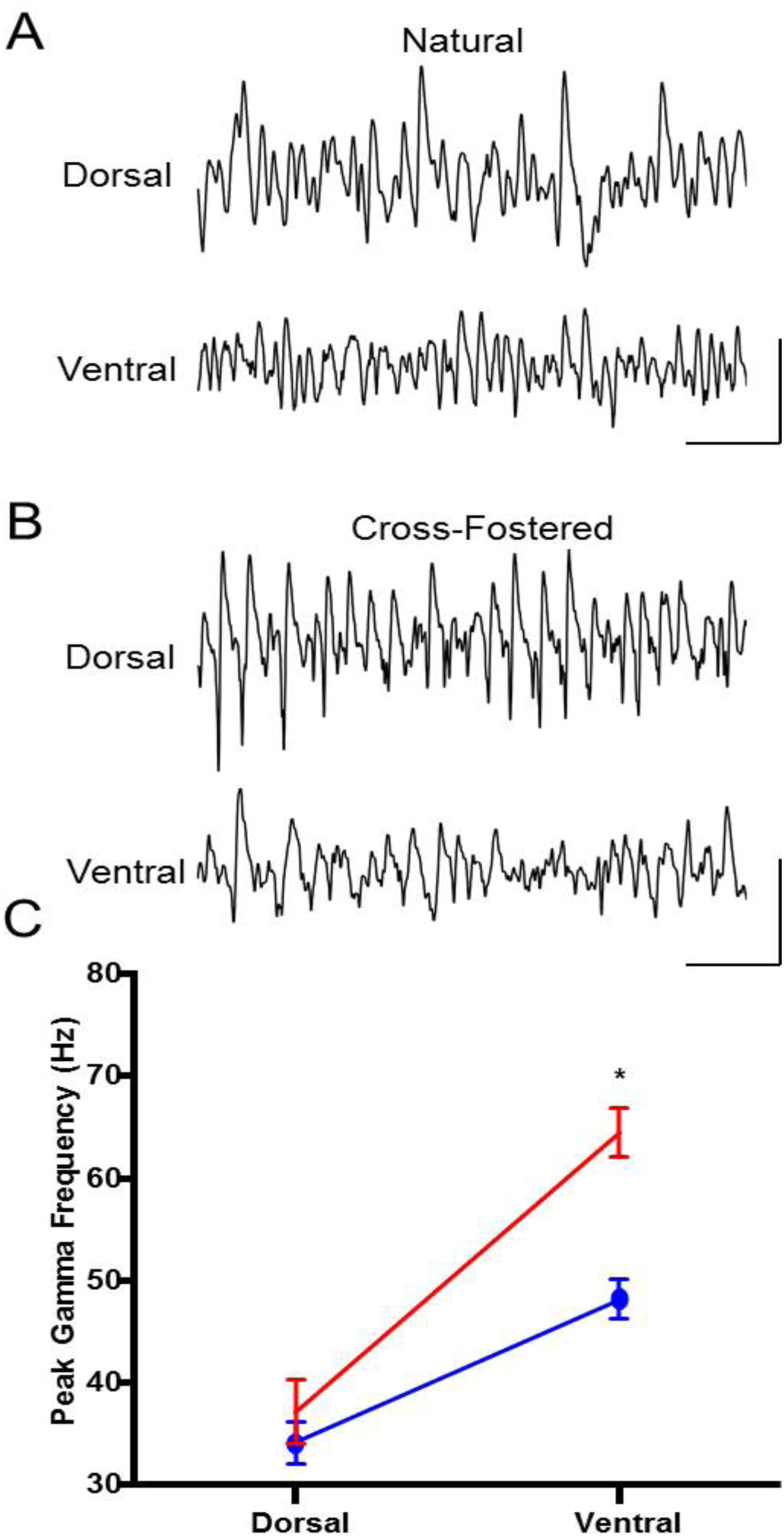
The dorsoventral gradient of the peak frequency of gamma oscillations is flattened in cross-fostered rodents. (A) Representative traces illustrate local field potential recordings obtained simultaneously from positions shown from the dorsal and ventral regions in natural slices. (B) Representative traces illustrate local field potential recordings obtained simultaneously from positions shown from the dorsal and ventral regions in CF slices. (C) Summary data for the peak frequency of gamma oscillations across the dorsoventral gradient. Scalebars: 100 μV, 0.1 s.

In slices from CF rats there was also a significant increase of the peak frequency across the dorsoventral axis (dorsal: 34.9 ± 2.9 Hz, ventral: 34.9 ± 2.9 Hz, n=26 slices, 10 animals; P<0.05, Fig 6C). When examining gamma power, a significant difference was observed between the dorsal (150.9 ± 12.1 μV^2^) and the ventral (16.8 ± 7.2 μV^2^) mEC regions (P<0.05, n=17 slices, 6 animals) data not illustrated. This difference was also maintained across the dorsoventral axis in slices from CF rats (dorsal: 87.7 ± 3.9 μV^2^, ventral: 21.3 ± 4.3 μV^2^, n=26 slices, 10 animals; P<0.05) data not illustrated.

## Discussion

The present data demonstrate that cross-fostering resulted in mild behavioural changes in both dam and pup. Once the pups matured, *in vitro* recordings revealed decreases in maximal gamma power evoked by kainate in both mEC and the CA3 subregion of the hippocampus. In addition, a large decrease in maximal frequency of gamma rhythms was seen in ventral mEC. These mEC gamma rhythm changes were accompanied by increased maximal GABA_A_ receptor-mediated IPSPs in both mEC stellate and pyramidal cells and an increase in parvalbumin expression selectively in LIII in this region. These findings were in contrast to those reported in severe early life stress models such as maternal deprivation, separation and pub handling-arguing, at least in part, against cross-fostering as an early life stress model.

However, CF rodents had a significantly reduced body weight throughout the 21-day weaning period when compared to N rodents and spent more time out of the nest. Perhaps in response to this, dams built more nests when caring for CF pups than their own biological progeny. Early life stress models have previously been shown to permanently decrease body weight but also to disrupt measures of maternal care such as LG or ABN^42–44^. No such maternal care deficits were seen with cross-fostering alone (Fig. 1).

From the above it seems that cross-fostering may generate some degree of stress in the pup but, unlike more overt early-life stress models, does not detrimentally affect the quantity and quality of maternal care given. From the seminal studies of Bowlby^4^ onwards it has been shown that adverse events in early life lead to an increased incidence of mental health problems in humans^45–48^ including, in severe cases, a link between foster care or institutionalization and varying symptoms of, or precursors to, psychiatric illness^10–12,49^. However, whether these observations reflect a negative influence of early life care by non-biological parents, or reflect the effects of stressors prior to this care is often unclear. The cross-fostering model used here may therefore help to disentangle the effects of these events (the situation leading to care by non-biological parents and the care itself) on brain function in adulthood.

Results from severe early life stress models point to selective changes in the parahippocampal region (see introduction). Communication within this region is afforded by gamma rhythms and slower rhythms (such as theta) that nest the gamma activity. These rhythms promote the transmission of spatial information^50–52^ and are vital for memory formation^53^. Post-institutionalized, adopted children show deficits in spatial span task performance when compared to controls^54^ and gamma/theta coupling has been shown to be significantly lower in institutionalized children^55^. In animal models of severe early life stress, deficits in gamma oscillations within the CA3 region of the hippocampus have been observed^56^ and gamma rhythmogenesis reveals a region-specific pattern of deficits in some animal models of schizophrenia^24,25^. However, data from patient studies and animal models resembling at least some of the genetic correlates of this psychiatric syndrome are far from equivocal (e.g. Hunt et al.,^57^). Consequently, the changes in gamma rhythm generation seen in the present study alone are hard to interpret in terms of potential mechanisms underlying the increased risk in psychiatric illness following early-life periods in care in humans.

A far more robust biological correlate of psychosis is the degree of expression of parvalbumin (PV) in interneurons^58,59^. A marked reduction in PV expression in post-mortem brain samples has led to arguments relating interneuron dysfunction to many symptoms of psychotic illness (e.g. Lewis et al., ^60,61^) and deficits in PV expression disrupt *in vitro* models of gamma rhythmogenesis^39^. Severe early life stress models have demonstrated decreases in parvalbumin positive interneurons in the prefrontal cortex^62^ (Holland et al., 2014) and the hippocampus^56^ that appear to correlate with increased incidence of psychosis in humans following time in care. In addition, decreases in both PV and calbindin expression has been seen observed in isolation reared rodents^63^.

In contrast, here we found an *increase* in PV expression (Fig. 5), strongly arguing against a correlation between foster care alone and increased psychosis, at least in this model. Such increases in PV expression have a precedent in some early life stress models. They have been linked to accelerated brain maturation^64^ through the early life ‘critical period’^65^ and perhaps reflect environmental experiences leading to elevated fear levels^66^.

Relating the observed increase in PV expression in LIII of mEC to the reduced gamma power and frequency seen in the present study is not straightforward. PV levels have been shown to have an exquisite developmental sensitivity to NMDAR hypofunction^67^ and here we show that the effects of cross fostering could be mimicked by the broad-spectrum NMDA receptor antagonist ketamine (Fig. 3). Changes in NMDA receptor function, specifically via GluN2A-containing receptors, has been shown to reduce PV levels^38^ but also concurrently alter GAD67 expression – an observation not seen here. In addition, some severe early life stress models have demonstrated a reduction in GluN2A- and GluN2B-containing NMDA receptors^68,69^, while others show a mechanistic link between elevated GluN2A and reduced PV expression^70^. Again, the present cross-fostering dataset failed to reveal any significant changes in the effect of antagonism of these two NMDA receptor subtypes (Fig. 3). Instead, the pharmacological data pointed to a selective change in GluN2C/D-containing receptors. It is interesting to note that, while poorly selective in general, ketamine has a preference for blocking these GluN2C/D-containing receptors^71^. In addition, while acute, high doses of ketamine mimic many symptoms of psychosis and schizophrenia in general, the drug, at lower doses, is also associated with antidepressant and anxiolytic effects^72,73^.

Interneurons in superficial layers of mEC retain a large proportion of excitation via NMDA receptors in adulthood^34^. Reduction in this type of excitation reduces IPSP magnitude and gamma frequency^32^ suggesting a direct causal link. Increased IPSP amplitude and slower decay kinetics can also cause frequency reductions^36^. However, here we saw only modest changes in these measures of electrically stimulated IPSPs (thus bypassing any changes in endogenous interneuronal excitation, Fig. 4). We suggest that the increase in maximal amplitude and concurrent decrease in decay time may therefore reflect an attempt at compensation for the apparent decrease in glutamate-mediated excitation via GluN2C/D-containing NMDA receptors.

None of the present observations from CF rats pointed to changes in the parahippocampal gyrus that could be related to the elevated rates of psychosis in adults having spent time in care as children. The main effects of cross fostering in mEC appear to be selectively seen in the ventral, rather than dorsal parts of this structure (Fig. 6). There are many differences at the biochemical and functional level along this axis: coupling between NMDA receptors and SK subtype of potassium channels is greater in ventral parahippocampal areas; Differences in HCN channel and M-current density are seen^74,75^; grid scale during exploration is larger in ventral areas. Of particular relevance to the data presented in this study, and to early life stress in general, is the observation that ventral parahippocampal regions seem to be selectively involved in social cognition and emotional responses to stress^41,76^.

In conclusion, the changes in gamma rhythms, NMDA function and PV expression seen in the present study may reflect the deficits in social cognition associated with foster care^77^. In contrast, these changes suggest that foster care may produce brain changes antagonistic to the elevated risk for psychosis and associated cognitive dysfunction following severe early life stress. The observations of beneficial effects of foster care on existing cognitive dysfunction^78^, the decrease in traits associated with psychopathy^11^ and general decrease in symptoms of psychosis^79^ may, we suggest, manifest in part from the changes in NMDA receptor-mediated drive to gamma-generating interneurons seen here.

## Materials and Methods

### Cross-fostering procedure

Procedures were performed in accordance with the UK Animals (Scientific Procedures) Act 1986 and the European Union Directive 2010/63/EU. Pregnant Wistar rats (Charles River, Kent, UK) were delivered at 10 days of gestation, and housed separately in modified clear polycarbonate boxes 480mm x 375mm x 210mm (Tecniplast, London, UK) containing aspen bedding for the duration of gestation and pre-weaning, except for during the cross-fostering procedure and cage cleaning. Dams were provided with nesting material. Food and water was provided ad libtium and temperature and humidity were controlled at 21±10C and 50% respectively. Animals were housed on a 12-hour light/dark cycle, lights on at 8:30 and off at 20:30. To enable viewing of maternal behaviours at night, red lights were switched on between 20:30 and 8:30. Pups were weighed on PND6, PND13 and PND20 with cage cleaning weekly commencing on PND6. Husbandry was performed at the same time of day by the experimenter in order to minimize any disruption to maternal behavior. An in-house cross-fostering procedure was performed on PND0. In the cross-fostered condition, two dams at a time were simultaneously separated from pups less than twelve hours after giving birth and placed in a holding cage out of sight. To ensure any effects on maternal care were a result of the cross-fostering procedure and not the physical effects of maternal separation and neonatal handling known to alter LG, a control procedure was carried out on N litters. This involved removing the dam from the home cage and placing in a holding cage out of sight of the pups. Briefly pups were handled for 3 seconds and placed back in the nest. In order to ensure that litter size was not a confounding factor in our experiments, litters were culled to limit the number of pups to eight in each litter and only males were used to negate against variations produced by hormonal fluctuations associated with the female reproductive cycle.

### Maternal behavior assessment

To permit recording of dam behaviour towards pups, a camera was suspended above each cage. Mirrors were positioned behind each cage to allow for optimal viewing and quantification of dams nursing postures. Videos were recorded on PND0-7, PND10, PND13 and PND17. Observational periods occurred in the dark phase of the light/dark cycle (22:00hrs and 07:00hrs), and in the light phase of the light/dark cycle (11:15hrs, 14:15hrs and 16:30hrs) to account for temporal fluctuations in arched back nursing (ABN) behaviours. Video recordings lasted twenty minutes and animals were undisturbed for the duration of recording. From these twenty minutes, twenty-five second epochs of video were analysed manually for behaviours of interest: at 3, 5, 7, 9, 11, 13, 15 and 17 minutes. Maternal behaviours of interest during the study were: the dam in contact with any number of pups, dam licking pups, dam arching over pups in an arched back nursing posture, dam passively nursing pups, number of nests and time pups spent out of nest.

### Electrophysiological experiments

Slices (450μm) containing hippocampus and mEC were prepared from young adult male naturally reared animals (N) or from cross-fostered (CF) rats. Animals were anesthetized with inhaled isoflurane, immediately followed by an intramuscular injection of ketamine (100 mg/kg) and xylazine (10 mg/kg). Animals were perfused intracardially with 50-100ml of modified artificial CSF (ACSF), which was composed of the following (in mM): 252 sucrose, 3 KCl, 1.25 NaH_2_PO_4_, 24 NaHCO_3_, 2 MgSO_4_, 2 CaCl_2_(H_2_O)_2_, and 10 glucose. All salts were obtained from BDH Chemicals (Poole, UK) or Sigma-Aldrich (Poole, UK). The brain was removed and submerged in cold (4–5°C) ACSF during dissection. Horizontal slices were cut and transferred to a recording chamber maintained at 34°C at the interface between ACSF [containing the following (in mM): 126 NaCl, 3 KCl, 1.25 NaH_2_PO_4_, 24 NaHCO_3_, 1 MgSO_4_, 1.2 CaCl_2_(H_2_O)_2_, and 10 glucose] and warm, moist carbogen gas (95% O_2_/5% CO_2_). Slices were permitted to equilibrate for 45 min before any recordings commenced. All slices (hippocampal and mEC) were taken from a ~2-mm-deep region containing mEC and temporal hippocampus (interaural, 2.56 – 4.36 mm; bregma, –7.44 to –5.64 mm). To examine neuronal oscillations across the dorso-ventral axis of the mEC, sagittal slices were prepared.

Recording electrodes were pulled from borosilicate glass (Harvard Apparatus, Edenbridge, UK), filled with ACSF, and had resistances in the range of 2–5 MΩ (extracellular) and 50–200 MΩ (intracellular). Neuronal subtypes were characterized electrophysiologically (interneurons having fast, non-accommodating spiking patterns, stellate cells having a pronounced sag on injection of hyperpolarizing current, pyramidal cells with an absence of this sag) and anatomically post hoc after biocytin injection. For stimulation experiments bipolar platinum electrodes (25µm diameter wire; tip separation, 200µm) were used to activate synaptic pathways electrically (1s duration; 1–10 mV). The stimulation electrode was positioned in the superficial layers of the mEC.

The following drugs were added to the ACSF for various experimental conditions: D,L-2-amino-5-phosphonovalerate (D-AP5; 50µM), (1*R*\*,2*S*\*)-*erythro*-2-(4-Benzylpiperidino)-1-(4-hydroxyphenyl)-1-propanol hemitartrate (Ifenprodil, 10µM), (2*S*\*,3*R*\*)-1-(Phenanthren-2-carbonyl)piperazine-2,3-dicarboxylic acid (PPDA, 100nM), 6-Imino-3-(4-methoxyphenyl)-1(6*H*)-pyridazinebutanoic acid hydrobromide (Gabazine, 1µM), (2*S*)-3-[[(1*S*)-1-(3,4-Dichlorophenyl)ethyl]amino-2-hydroxypropyl](phenylmethyl)phosphinic acid hydrochloride (CGP55845, 1 µM), Carbamoylcholine chloride (carbachol) and 2,3-Dioxo-6-nitro-1,2,3,4-tetrahydrobenzo[*f*]quinoxaline-7-sulfonamid (NBQX, 10 µM) from Tocris Bioscience (Avonmouth, UK), Kainic acid (1-500nM) and ketamine (25µM) from Sigma–Aldrich (Poole, UK), NVP (50nM) from Merck, UK).

Peak frequency and power values were obtained from power spectra of 60 second epochs of data and generated with Fourier analysis in the Axograph software package. Monosynaptic IPSP amplitude was determined in the presence of blockers for GABA_A_ (Gabazine), AMPA (SYM 2206), Kainate (NBQX) and NMDA (NBQX). Cells were characterised and depolarized to approximately −20- −30mV in order to observe IPSPs. A bipolar stimulating electrode was placed in the same cortical layer as the cell and increasing voltage stimulations were applied to the slice using an isolated stimulator box. The amplitude of IPSPs was determined by calculating the difference between the resting membrane potential before the stimulation and the peak of the IPSP averaged from 20 stimulations at each intensity.

### Immunocytochemical experiments

Slices were maintained at 4°C for >24 hrs. in 4% Paraformaldehyde (PFA) in 0.1M phosphate buffer solution (PBS). For cell count experiments, slices were then embedded in a 10% solution of gelatine (type A Porcine; Sigma-Aldrich) in 0.1M PBS, which was then dissolved at 40°C for 20 minutes. Slices were submerged in gelatine and left to set at 4°C for 2hrs before being removed and stored in PFA at 4°C overnight. Gelatine-containing slices were subsequently removed from the fixative and underwent three 10-minute washes in 0.1M phosphate buffer (PB) before sectioning. Slices were re-sectioned to 60μm to ensure successful visualisation of immunohistochemically-stained cells Within cell counting protocols; slices were washed in 10mM sodium citrate for 1 hour at 80°C to denature any residual proteins and bleached in 0.1% gelatine, 0.1% triton and PBS for at least 1 hour.

Slices were then incubated overnight with the primary antibody of either mouse parvalbumin (Millipore, UK), mouse GAD67 (Millipore, UK), mouse calretinin (Millipore, UK) or somatostatin (Millipore, UK) at a ratio of 1:5000, whilst controls were incubated with w/o antibody. After overnight incubation, slices were washed in PBS then incubated in GAM Biotin (Millipore, UK) for 2 hours at room temperature. They were finally incubated with the ABC elite kit (Vector Labs, Peterborough) then washed in PBS then mounted on slides with Mowiol. A single individual performed quantifications, independently, and blind to experimental conditions and to the origin of the sections from N reared or CF animals. Cells were considered immunopositive when they showed a heavy brown labeling corresponding to the reaction product. Only positive cells were counted in microscopic view fields at 10x magnification within the 60μm sections of the mEC and hippocampus. The counts were then scaled to fit 450μM sections using dimensions regarding the cortex described in Du et al, 1995.

### Statistical analysis

Normality of the data was tested using the Shapiro-Wilk normality test and can be assumed to pass the normality test unless otherwise stated. For parametric data, paired t-tests were used to measure any differences between the means of data before and after a manipulation from the same slice or cell. Alternatively, if data was non-parametric a Mann-Whitney test was used. For concentration curve analysis a 2-way, repeated measures Analysis of Variance (rmANOVA) test was used with compensation for multiple comparisons using the Holm-Sidak method (Sigmastat, Ststat inc. San Jose, CA).

## Acknowledgements

This work was supported by The Wolfson Foundation, The Royal Society and the Newcastle upon Tyne Healthcare Charities Trust. SH was funded by a CASE studentship.

## Author Contributions

S.H., C.H.D., M.A.W. and M.O.C. designed research; S.H. K.H., G.L., T.A., A.S., and M.O.C. performed research; S.H., K.H., G.L., T.A., A.S., M.A.W. and M.O.C performed analysis of data; and S.H., M.A.W. and M.O.C. wrote the manuscript.

The authors declare no conflict of interest.

